# Predicting efficiency of writing short sequences into the genome using prime editing

**DOI:** 10.1101/2021.11.10.468024

**Authors:** Jonas Koeppel, Elin Madli Peets, Juliane Weller, Ananth Pallaseni, Fabio Liberante, Leopold Parts

## Abstract

Short sequences can be precisely written into a selected genomic target using prime editing. This ability facilitates protein tagging, correction of pathogenic deletions, and many other exciting applications. However, it remains unclear what types of sequences prime editors can easily insert, and how to choose optimal reagents for a desired outcome. To characterize features that influence insertion efficiency, we designed a library of 2,666 sequences up to 69 nt in length and measured the frequency of their insertion into four genomic sites in three human cell lines, using different prime editor systems. We discover that insertion sequence length, nucleotide composition and secondary structure all affect insertion rates, and that mismatch repair proficiency is a strong determinant for the shortest insertions. Combining the sequence and repair features into a machine learning model, we can predict insertion frequency for new sequences with R = 0.69. The tools we provide allow users to choose optimal constructs for DNA insertion using prime editing.

## INTRODUCTION

Efficient insertion of short DNA sequences into genomes could change the course of biotechnology and medicine. Small insertions can encode protein tags for purification and visualization, or allow manipulation of protein localization, half-life, and interaction profiles to control their function. Integrating sequences for transcription factor binding sites and splicing modulators provides control over gene expression, while introducing structural elements or recombinase sites can change DNA conformation and provide a substrate for large-scale engineering^1,2^. For therapeutic opportunities, over 16,000 small deletion variants have been causally linked to disease^3,4^, and could in principle be restored by inserting the missing sequence^5,6^. A prominent example is cystic fibrosis, where 70% of cases are caused by a 3 nt deletion^7,8^. An ideal tool to enable these applications would integrate the edits efficiently, accurately, and safely, avoiding unintended outcomes or double strand break stress that has hampered Cas9-based therapies^9-11^.

Prime editors can insert short DNA sequences without generating double-strand breaks or needing an external template. They consist of a nicking version of Cas9 fused to a reverse transcriptase domain, which is complexed with a prime editing guide RNA (pegRNA)^12^. The pegRNA comprises a primer binding site homologous to the sequence in the target, and a reverse transcriptase template that includes the intended edit, all in the 3’ extension of a standard CRISPR/Cas9 guide RNA. At the target site, Cas9 nicks one strand of the genomic DNA, which then anneals to the primer binding site on the pegRNA, and is extended by the Cas9-fused reverse transcriptase using the pegRNA-encoded template sequence. Next, DNA repair mechanisms resolve the conflicting sequences on the two DNA strands, ultimately writing the intended edit into the genome. When CRISPR/Cas9 was compared to molecular scissors capable of disrupting target genes, and base editors were called molecular pencils for their ability to substitute single nucleotides, prime editors were described as molecular word processors able to perform search and replace operations directly on the genome^13^.

The prime editing system is complex, and the determinants of its efficiency are not fully understood. Several partly independent steps, including three DNA binding events and a successful mismatch repair are needed to produce an edit, each potentially introducing biases. In the largest study so far to understand these biases, Kim et al. comprehensively tested the consequences of varying the reverse transcription templates and primer binding site lengths using a library of 55,000 pegRNAs. Editing rate increased with Cas9 gRNA activity, as well as GC content and melting temperature of the primer binding site. Primer binding sites of 13 nt and reverse transcriptase templates of 12 nt generally worked well^14^.

The majority of libraries used by Kim *et al*. contained the same single nucleotide substitution 5 nt upstream of the nick site. Similarly, nearly all current characterization of prime editing efficacy has predominantly focused on single nucleotide substitutions^12,15-18^. Of the many possible useful sequences in molecular biology, only a handful have been introduced with prime editing and the longest successfully reported insertion was 44 nt in length^12^. Therefore, in contrast to relatively deep understanding of Cas9 mutagenesis^10,19-21^ and base editing outcomes^22-24^ very little is known about how the inserted sequence affects efficiency, and the length range of insertions feasible by prime editing has not been defined.

Here, we systematically measure the insertion efficiency of over 2,600 sequences and identify the features responsible. We find that insertion sequence length, nucleotide composition, secondary structure, and repair pathway activity together explain most of the variation in insertion rate. We then use these insights to train a sequence-based prediction model informed by mismatch repair efficiency that predicts editing outcomes for novel sequences with high accuracy, and allows selection of optimal reagents for new insertions.

## RESULTS

We sought to systematically characterize how the length and composition of inserted sequence, as well as cell line, target site, and the version of the prime editor system affect insertion rates. To do so, we designed 2,666 pegRNAs encoding insertions immediately upstream of the nick site. These comprise 270 sequences useful for molecular biology (including e.g. His-6 tag, recombinase sites, and mNeonGreen11^25^), 1,957 eukaryotic linear motifs^26,27^, and 439 sequences with variable secondary structure (Figure 1a). The insertion lengths ranged from 1 to 69 nt, with varied GC content (Figure 1b). We used lentiviruses to deliver the library against four target sites (three previously tested: HEK3, EMX1, FANCF^12^ and the safe-harbor CLYBL locus^28^) in two cell lines (HEK293T and HAP1), followed by transient transfection of the prime editor plasmid, five days of selection, and sequencing of two amplicons from the cell pool, one of the targeted locus and one of the pegRNA locus (Figure 1c). We calculated insertion efficiencies as the fraction of reads in the target site amplicon with a given insertion divided by the fraction of reads for the pegRNA encoding it in the pegRNA amplicon, and analyse them as the main statistic in the rest of the study.

**Figure 1.**
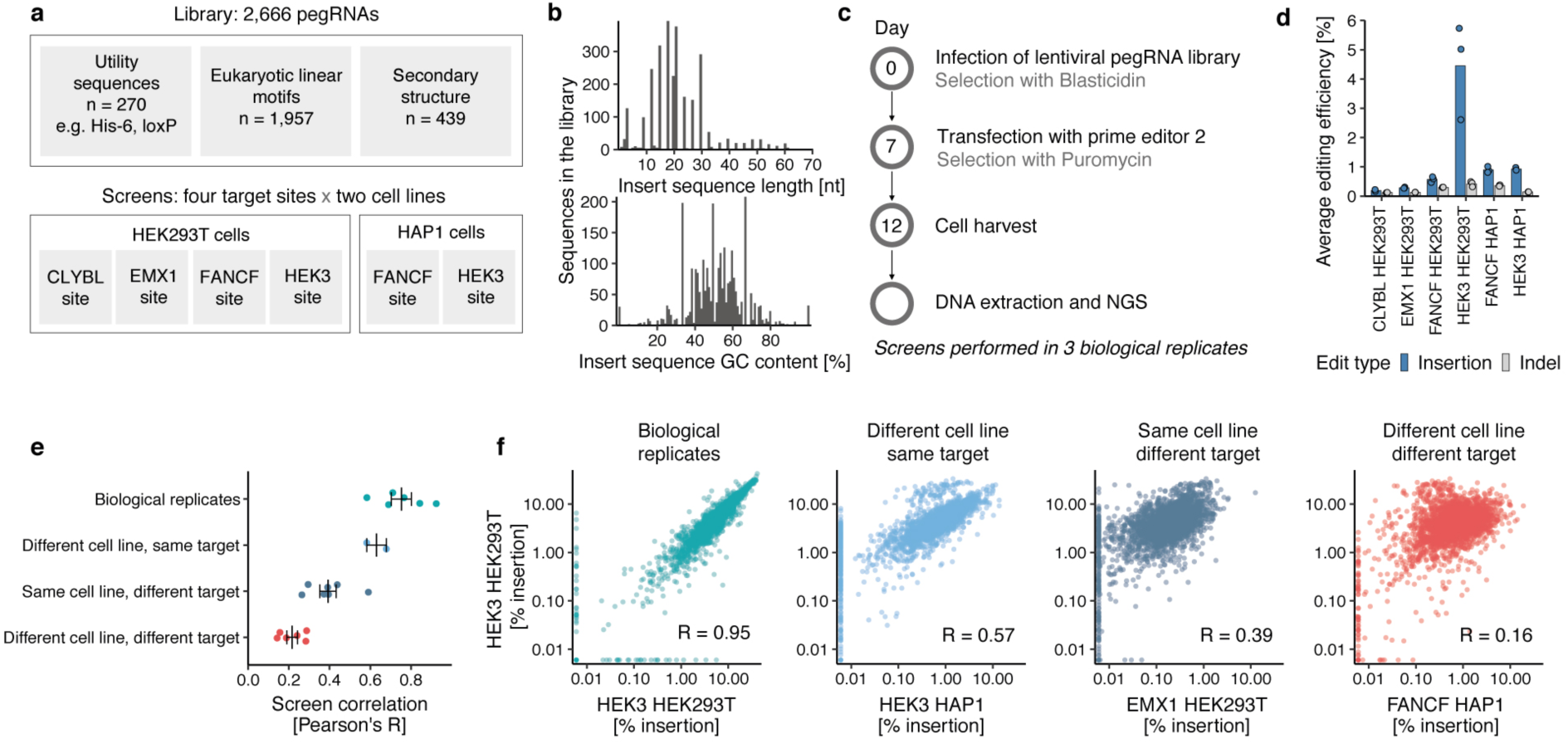
High-throughput measurement of prime insertion efficiencies. **a**. Screen setup. **b**. Library composition. Number of sequences in the library (y-axis) with different insert sequence lengths (x-axis, top panel) and %GC content (x-axis, bottom panel). **c**. Experimental design **d**. Editing frequencies. Average mutation frequency (y-axis) for different screens (x-axis) stratified by mutation type (blue: insertions; grey: indels). Markers represent one replicate and bars the average across *n*=3 biological replicates. **e**. Replicate concordance. Pearson’s R between insertion rates in two screens (x-axis) for different comparisons (y-axis, colors). Markers: correlation value of one pair of screens (for replicate correlations, mean of pairwise comparison across *n*=3 biological replicates); line and whiskers: mean and standard error of mean. **f**. Representative examples of categories from (e). Percent insertion in the HEK3 locus in HEK293T cells (y-axis) compared to values (x-axis) in other contexts (panels, colors) for insertion sequences (markers). Left panel: comparison of biological replicates; other panels: comparison of replicate averages. Label: Pearson’s R of values in linear scale. Colors: as in (e).

Insertion efficiencies of pegRNAs varied widely. The top 1% of pegRNAs were inserted 210-2,040 times more efficiently than the bottom 1% across the various target site and cell line combinations (Supplementary Figure 1a), indicating substantial sequence-dependent variation. The insertion rates were highly consistent across biological replicates (average Pearson’s R=0.76, range 0.64-0.95; Supplementary Figure 1b), but differed in magnitude across screens (average across pegRNAs 0.19% for CLYBL locus in HEK293T to 4.45% for HEK3 locus in HEK293T cells, Figure 1d). Small insertions and deletions around the target site were rare (0.12%-0.42%, Figure 1d), as were additional single-nucleotide mutations around the nicking site derived from the prime editing process (0.03% on average in reads with an inserted sequence vs 0.03% in reads without insertions, Supplementary Figure 1c). Overall, the intended insertions were the dominant mutations generated, and we do not consider the unintended edits further.

To understand the consistency of insertion efficiencies across contexts, we next compared them between replicates, cell lines, and target sites. Insertion rates into the same target site in different cell lines were more correlated (mean Pearson’s R=0.63) than into different target sites in the same line (mean Pearson’s R=0.39), indicating a greater dependence on target locus than cellular background. The correlation was weakest when both target site and cell line were different (mean Pearson’s R=0.22, Figure 1e-f), demonstrating both target sequence-specific, and cell line-dependent biases on insertion.

### Insert size effects

Given the repeatable sequence dependent variation in insertion rates that spans over three orders of magnitude, we sought to understand the responsible features. We first asked how insert sequence length affects insertion efficiency. For HEK293T cells, sequences of up to 4 nt (all four single nucleotides, all 16 dinucleotides, and 63 trinucleotides) were inserted on average 2.1-4.6 times more efficiently than longer ones across the four targeted sites (Figure 2a). Insertion frequency did not decrease monotonically with increasing insert length, as sequences between 15 and 21 nt were inserted 1.3-1.6 times more efficiently than 10-14 nt ones (Figure 2a). These relative biases in efficiency were shared between all target sites, despite a 20-fold range of their average insertion rates. Inserts longer than 45 nt were incorporated less frequently, albeit at a screen average rate of 22-38% of sequences shorter than 45 nt. The longest sequence that was inserted at > 1% frequency (1.4%, HEK3 site in HEK293T cells) was 66 nt, demonstrating that reasonably efficient integration of moderately long sequences is feasible with prime editing.

**Figure 2.**
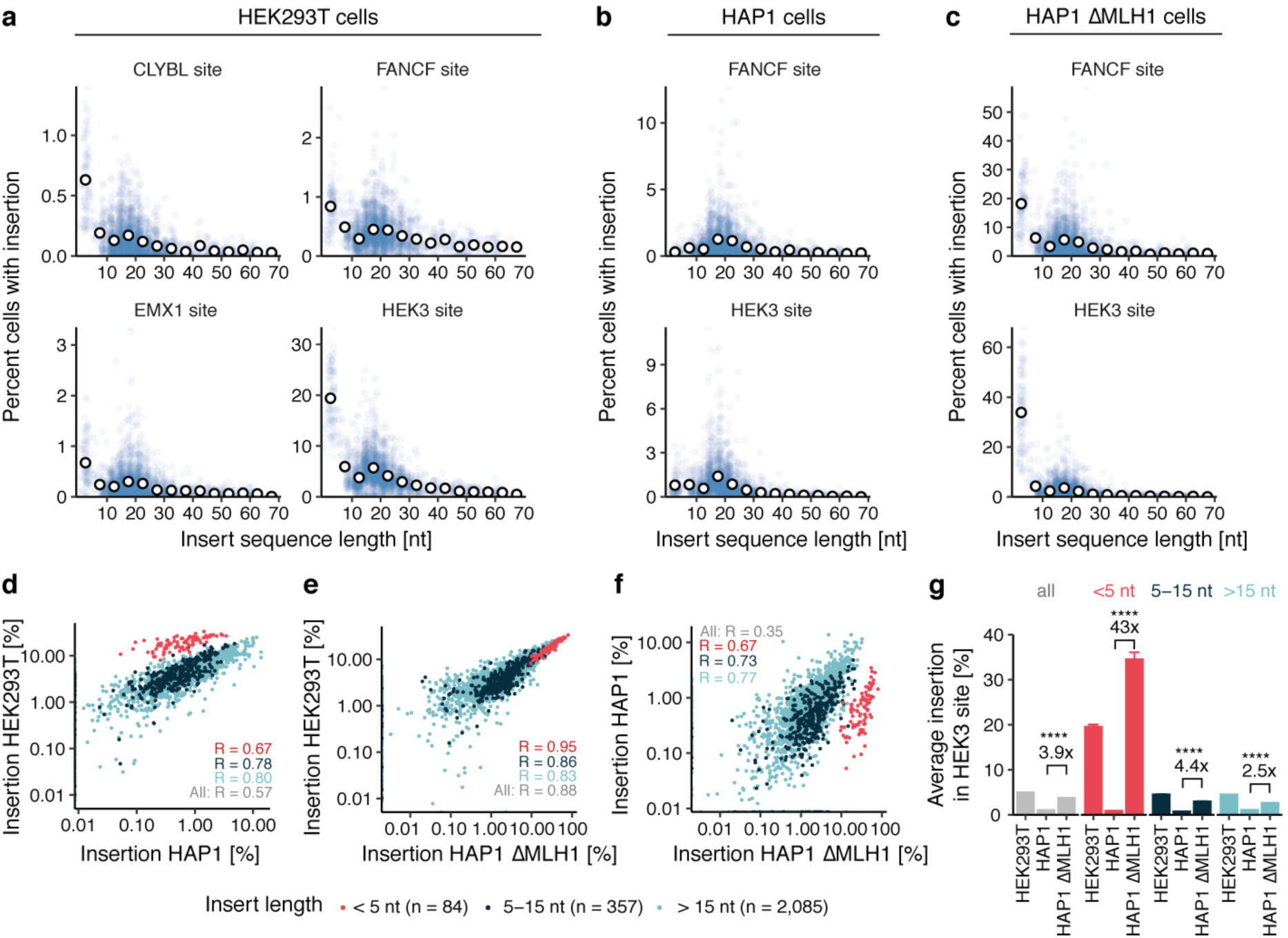
Prime insertion efficiency depends on insert length. **a**. Insertion rate in HEK293T cells. Percent cells with insertion (y-axis) for different insert sizes (x-axis) of individual sequences (blue markers) and their averages in 5nt length bins (white markers) at different target sites (panels). Data represent the average of *n*=3 biological replicates. **b**. As a), but for HAP1 cells. **c**. As a), but for HAP1 ΔMLH1 cells. **d**. Insertion rate in HEK293T cells (y-axis) compared to rate in HAP1 cells (x-axis) at the HEK3 target of individual sequences (markers). Red: short sequences (up to 4nt); blue: medium sequences (5-15nt); teal: longer sequences (>15nt). Label: Pearson’s R between rates. The data are an average from *n*=3 biological replicates. **e-f)**. As (d), but comparing insertion rates in HAP1 ΔMLH1 cells with HEK293T cells (e) and HAP1 wild type cells (f). **g**. MLH1 knockout increases insertion rates disproportionately for short sequences. Average insertion rates at the HEK3 locus (y-axis) across different cellular contexts (x-axis) for different length bins (colors). Comparison: ratio of average insertion rate between HAP1 ΔMLH1 cells and HAP1 wild type cells; p-values (all < 1×10^−33^) from two-sided Student’s t-test. Error bars: standard error of the mean. Colors as in (d)-(f).

In contrast to HEK293T cells, insertion frequency of the short 1-4 nt sequences was 1.2 to 3.7-fold lower than that for longer ones in HAP1 cells (Figure 2b). This reduced the correlation of insertion rates between the two cell lines at the same site compared to replicate correlation (Pearson’s R=0.68 for FANCF and 0.57 for HEK3, Figure 2b; median replicate correlation R=0.75, Supplementary Figure 2a). However, stratifying the inserted sequences by length recovers strong concordance, with correlations between rates in HAP1 and HEK293 cells at the HEK3 locus increasing to 0.67, 0.78, and 0.80, respectively, for sequences of lengths 1-4 nt, 5-15 nt, and 16-70 nt. This correlation within length bins indicates consistent pegRNA effects, while the discordance between bin averages across lines (but not targets in the same line) hints that the cellular context does not influence inserts of different lengths in the same way. One possible explanation is mismatch repair (MMR) proficiency, since HEK293T cells are partly MMR deficient due to promoter methylation of MLH1^29^, while HAP1 cells are not. The MMR pathway recognizes and excises short mismatches of less than 13 nt and could therefore remove short insertions in HAP1 cells before the nicked strand is re-ligated^17,30^.

To test the hypothesis that rates of inserting short sequences differ between cell lines due to mismatch repair activity, we screened the HEK3 and FANCF-targeted libraries in HAP1 cells that are knockout for MLH1 (HAP1 ΔMLH1, Supplementary Figure 2b,c). We found that average insertion rates increased 3.9 to 5.1-fold in the mutant background compared to wild type HAP1 cells. The rates 1-4 nt sequences were most affected, increasing by 43-66 fold to 19-34%, while the rates of 5-15 nt and >15 nt sequences increased 4.4-7.9 fold and 2.5-4.2 fold, respectively (Figure 2g, Supplementary Figure 2d,e). These remarkable increases in insertion rates for short sequences are consistent with a model where MMR predominantly recognizes short insertions, and thereby antagonizes prime editing. Indeed, for the HEK3 target site, insertion rates in HAP1 ΔMLH1 cells are as correlated to HEK293T cells as replicates (Pearson’s R = 0.88, Figure 2e, Supplementary Figure 1b) while correlations between wild type HAP1 and HAP1 ΔMLH1 cells are modest (Pearson’s R = 0.35, Figure 2f) and improve when stratifying by sequence length. The same pattern of higher insertion rates for short sequences in ΔMLH1 cells was observed for the FANCF locus (Supplementary Figure 2f,g). These findings highlight that MMR proficiency is the major source of variation between the tested cellular contexts for prime insertions.

### Sequence effects

We next examined the length-independent causes of variation in insertion rate. We calculated the relative insertion rate for each insert by dividing its marginal rate by the median in a 5-nt length bin (Methods), and observed it is positively correlated with GC content across all target sites and cell lines (Figure 3a). Each extra percentage in GC content increased the relative insertion rate between 1.8% (FANCF site in HEK293 cells) and 4.9% (EMX1 site HEK293T cells) on average. Specifically, we observed a strong cytosine preference for the CLYBL, EMX1, and HEK3 loci (each extra percent cytosine increases relative insertion rate by 2.7-5.4%), while both the percent of cytosines (0.9-1.2%) and guanines (1.3-3.3%) increase insertion rates for the FANCF locus. Conversely, percent of adenine and thymine decreased insertion rates for all loci and cell lines (−0.8% to -5.1% and -0.5% to -3.6% respectively, Supplementary Figure 3a,b).

**Figure 3.**
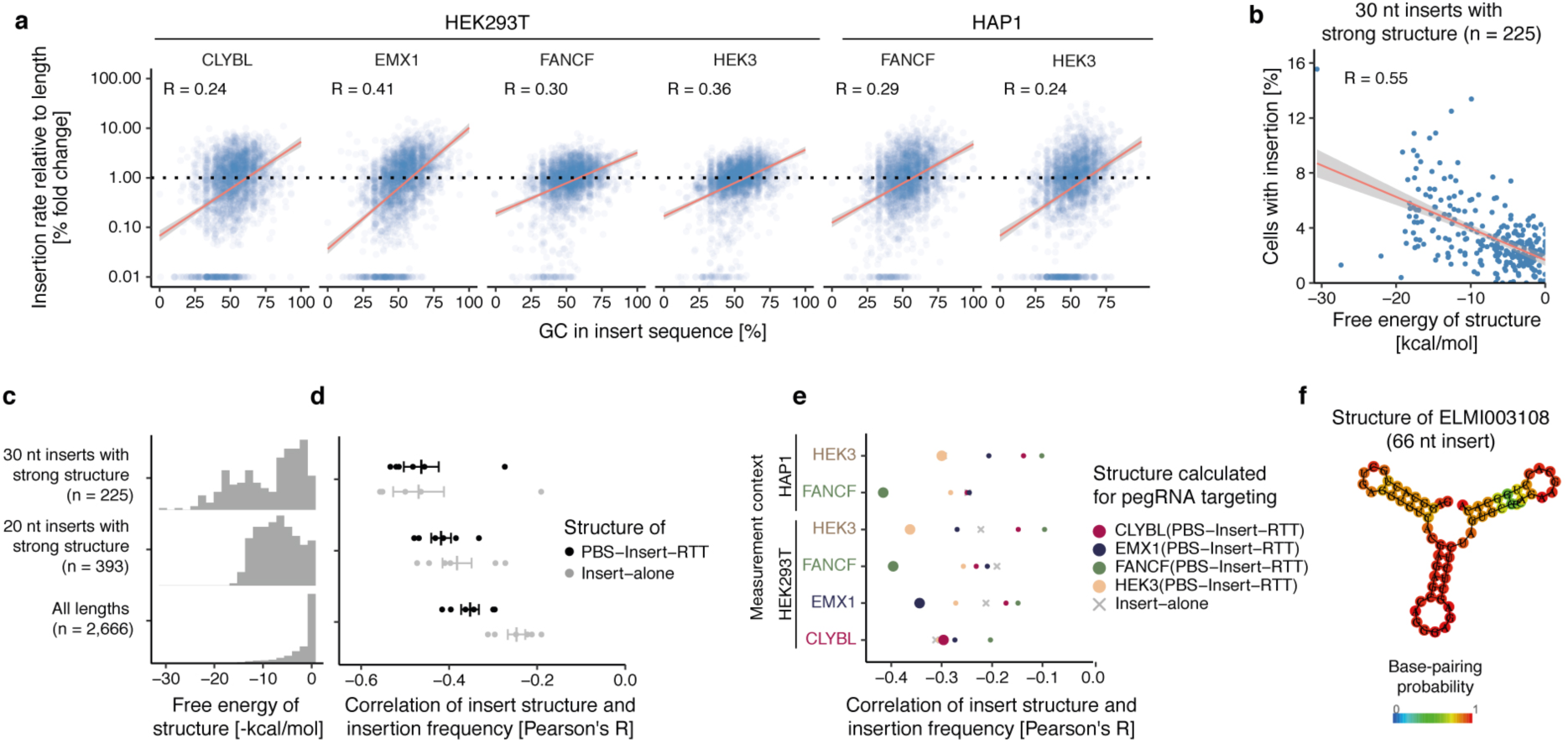
GC content and secondary structure of the insert sequence are positively correlated with insertion rate. **a**. Insertion efficiency dependence on GC content. Insertion rate relative to length bin average (y-axis) for inserts with different GC content (x-axis) of individual sequences (markers) at different target sites and cell lines (panels). Red line: linear regression fit; shaded area: 95% posterior confidence interval of the fit. Data represent the average of *n*=3 biological replicates. **b**. Insert sequence free energy correlation with insertion rate. Percent of cells with insertion (y-axis) for 30 nt sequences in the HEK3 locus in HEK293T cells (markers) with calculated Gibbs free energy (ΔG) from ViennaFold (x-axis). Red line: linear regression fit; shaded area: 95% posterior confidence interval of the fit. Data represent the average of *n*=3 biological replicates. **c**. Subsets of insertion sequences with fixed length and variable secondary structure. Frequency (y-axis) of insert sequence free energy (ΔG, x-axis) for 30 nt (top panel) and 20 nt sequences (middle panel), and the entire library (bottom panel). **d**. Correlation of insertion frequency and insert structure ΔG (x-axis) for 30 and 20 nt sequences, and the entire library (y-axis), stratified by whether ΔG was calculated for the insert sequence alone (grey) or for the entire 3’-extension, consisting of primer binding site (PBS), insert, and reverse transcription template (RTT; black) for different combinations of target sites and cell lines (markers). Data represents averages from *n*=3 biological replicates. Bars: mean and standard error of the mean. **e**. Target dependence of pegRNA secondary structure free energy correlation to insertion rate. Correlation (x-axis) between insertion efficiency measured in different target sites and cell lines (y-axis) and pegRNA 3’ extension structure free energy calculated for pegRNAs against different target sites (colored markers), as well as insert sequence alone (grey cross). **f**. Example long sequence with successful insertions. The predicted secondary structure of a 66 nt insert sequence (ELMI003108) which was inserted with 1.4% efficiency into the HEK3 locus in HEK293T cells. Colors: base pairing probability.

Our library contained 225 and 291 insert sequences of 20 nt and 30 nt respectively, that were designed to form secondary structures of varying strength, including some sequences with perfect hairpins. Their secondary structure free energy as quantified by the Vienna fold ΔG (more negative ΔG values indicate stronger secondary structures)^31,32^ was negatively correlated with insertion efficiency (average Pearson’s R = -0.38 and -0.50 for 20 nt and 30 nt insertions respectively, Figure 3b-d, Supplementary Figure 3c). Even when excluding the shortest inserts of up to 3 nt, most of the inserted sequences in the library do not form strong structures (Figure 3c) and the Vienna fold free energy explains less signal variation (average Pearson’s R=-0.25). We speculated that more important than the structure of the insert sequence alone is the structure across the entire pegRNA 3’ extension, comprising the primer binding site and the reverse transcription template, and indeed, more variation was explained when including the extension (average Pearson’s R=-0.35, Figure 3d, Supplementary Figure 3d-f). Since the extension is specific to the target, this partly explains the differences in insertion rates we observed across target sites. To quantify this effect, we measured how well free energies of pegRNA extensions designed for one target site predict insertion efficiency at another target site. Consistent with a target site-specific effect, the correlation was strongest when the target site was matching to the 3’ extension the structure it was calculated from (Figure 3e).

Combining insert sequence length, GC content and structure explained why some sequences inserted much better than others, which can help guide the choice of tags to insert. For example, the long 66 nt ELMI003108 sequence that was inserted in the HEK3 locus at 1.39% insertion frequency (0.66% on average for the other 10 sequences > 66 nt) had a GC content of 62% and formed a strong structure alone as well as within the pegRNA context (ViennaFold free energy = -23.8 and -37.5 respectively, Figure 3f). Other longer sequences that inserted well relative to their size were recombinase sites, presumably due to their secondary structure that often contains hairpins (Supplementary Figure 4a,b).

### CRISPR system effects

Finally, we considered how aspects of the CRISPR/Cas system itself impact insertion rates. First, it is known that the occurrence of four consecutive thymines acts as a transcription terminator for RNA polymerase III and strongly impairs guide RNA expression^33,34^. We confirmed that the average insertion rate for sequences that contain this tetranucleotide was 4.2 to 11.6-fold lower compared to sequences without (Figure 5a), while stretches of four adenines showed a weaker but significant effect (average 1.5 to 1.8-fold reduction, Supplementary Figure 4c). Overall, 21 of the 24 pegRNAs that were not inserted in any screen contained at least one instance of the TTTT sequence.

Second, to disentangle the contribution of the reverse transcription step, we made a prime editor construct with the nicking Cas9 fused to an engineered feline leukemia virus reverse transcriptase (MashUp RT - pipettejockey.com) with a similar fidelity to the murine leukemia virus RT used in PE2. The average insertion rates observed using this construct were 4.6-fold lower compared to the standard PE2 (0.98% and 4.56% respectively; Figure 4b), but as correlated to PE2 as another biological replicate (Pearson’s R=0.80; Figure 4c, Figure 1e), and with consistent marginal effects of the contributing features (Supplementary Figure 5). Therefore, the reverse transcriptase used is not a major cause of variation in insertion rate in our study.

**Figure 4.**
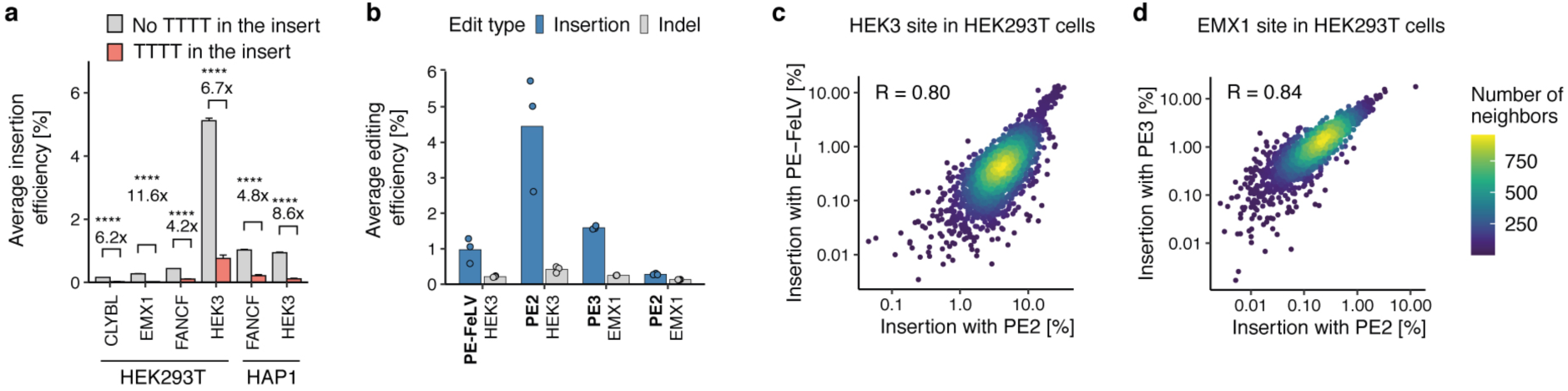
CRISPR system effects. **a**. Impact of four consecutive thymines. Average insertion rate (y-axis) for different screens (x-axis) stratified by presence of four consecutive thymines (grey: absent; red: present). Comparison: ratio of grey to red bar height. p-values derived from a two-sided student’s t-test with all p-values < 1×10^−67^. *n*=3 biological replicates. **b**. Editing frequencies for alternative prime editing systems. Mutation frequency (y-axis) for three biological replicate screens (markers) using different prime editor systems (x-axis) stratified by mutation type (blue: insertions; grey: indels). Bar: average of markers. **c**. Impact of an alternative reverse transcriptase. Insertion frequencies at the HEK3 site in HEK293T using the standard MMLV reverse transcriptase (PE2, x-axis) and the FeLV reverse transcriptase (PE-FeLV, y-axis) for different insertion sequences (markers). Colors: number of neighboring data points. Label: Pearson’s R. *n*=3 biological replicates **d**. As c), but comparing PE3, and at EMX1 site.

**Figure 5.**
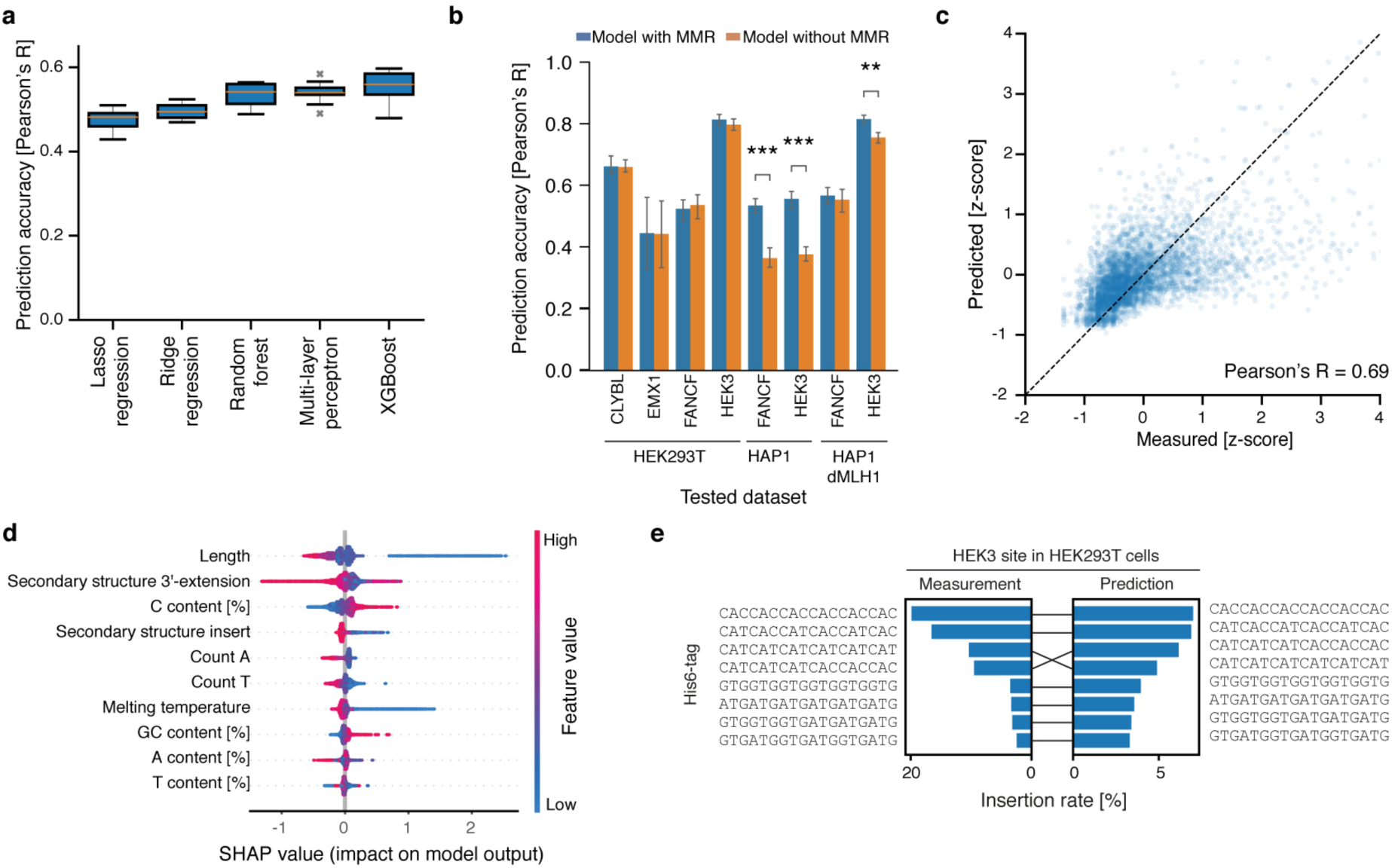
Predicting prime insertion efficiencies. **a**. Cross-validation performance of different models. Pearson’s R between predicted and measured held-out pegRNAs (y-axis) for ten cross-validation folds across a range of models (x-axis). Box: median and quartiles; whiskers: 90th percentile; cross: outlier. **b**. Impact of sequence-independent mismatch repair (MMR) proficiency feature. Pearson’s R between measured and predicted insertion rate on held-out pegRNAs (y-axis) on different screens (x-axis) for XGBoost model that includes MMR feature (blue) or not (orange). Whiskers: standard error of mean. Comparison: Student’s t-test between models with MMR and without; ***: p < 10^−3^; **: p < 10^−2^. **c**. Concordance of prediction and measurement. Predicted (x-axis) and measured (y-axis) insertion efficiency on the combined dataset for held-out pegRNAs (markers). Label: Pearson’s R. Dashed line: y=x. **d**. Feature importance. Distribution (row y-axis) of SHAP values (x-axis) for features ordered top to bottom by median absolute SHAP value (y-axis). Colors: feature value. **e**. His6-tag insertion rate. Insertion rate (x-axis) for Predicted (right) vs. measured (left) insertion rates of different alternative His6-tags (y-axis) into HEK3 locus in HEK293T cells

Including an additional sgRNA to nick the non-edited strand increases editing efficiency as well as indel formation rate^12^. We next explored how the addition of this extra sgRNA affects the insertion frequencies of our library. We chose the EMX1 locus in HEK293T cells where we observed poor insertion efficiencies of 0.28% on average without the nicking guide RNA, and co-transfected with a nicking guide RNA that targets 77 nt downstream of the pegRNA target^35^. We found that the extra nick increased the average insertion rate by 5.7-fold to 1.59%, and moderately increased the indel rate by 1.7-fold to 0.22% (Figure 4b). Importantly, the relative insertion rates for sequences in the library remained similar, with correlations in line with those of biological replicates and the other reverse transcriptase (Pearson’s R=0.84, Figure 4d), and consistent feature impact (Supplementary Figure 5) suggesting the determinants of prime insertion efficiency we uncovered for PE2 are also valid for PE3.

### Predicting insertion rates

Given our improved understanding of prime insertion efficiency, we next created a prediction model. We extracted salient features such as insert length, nucleotide composition, and folding energy for each pegRNA, and trained regression models of insertion rates. We chose to use XGBoost (gradient boosted decision trees^36^), as it achieved the best cross-validation performance (Figure 5a), and further evaluated its accuracy on datasets narrowed to individual sites and cell lines (Supplementary Figure 6a). The model generalized well (Supplementary Figure 6b), but we noticed that training on data only from HAP1 experiments resulted in worse performance on the HEK293T and HAP1 ΔMLH1 lines (average Pearson’s R = 0.31 vs 0.49 and 0.50 respectively). Given the observations of mismatch repair importance above, and the lack of repair proficiency as a feature input to the model, we hypothesized that including an experiment-specific and sequence-independent feature to capture this will improve prediction accuracy. Indeed, the model improved substantially when adding MMR status as a feature (increase in Pearson’s R > 0.1; paired t-test p < 10^−3^; Figure 5b).

Our final model is trained on data from experiments on all cell lines and target sites, and includes sequence characteristics as well as repair proficiency as features. It predicted insertion rates for unseen insert sequences with high accuracy (Pearson’s R=0.69, Figure 5c), which is very near the limit set by the correlation of biological replicates (average Pearson’s R=0.76). The features important for prediction^37^ reflected the observations above, with insert sequence length, secondary structure of the pegRNA, and nucleotide composition having the largest impact (Figure 5d). We call this method MinsePIE (Modelling insertion efficiency for Prime Insertion Experiments) and make it available at https://github.com/julianeweller/MinsePIE.

Predictive models inform experiments and help in selecting the components of a desired insertion. A common application of small sequence insertion is endogenous protein tagging, and the His6-tag is frequently used to enable purification. The possible tags that give equivalent amino acid sequences range in codon choice, thus resulting in pegRNAs with varying secondary structure and nucleotide composition when inserted. Our library contained 8 codon variations of the His6-tag in forward and reverse orientations. The average insertion difference between the best codon variant and the worst was 13.3-fold, highlighting the importance of choosing the optimal codon variant for insertion. We withheld the data for all His6-tags in our library from the training set, and predicted their efficacy of insertion into the HEK3 locus in HEK293T cells. The versions of the His6-tag that were both predicted and measured to have the highest insertion rates had the highest cytosine content by using mainly the CAC histidine codon (Figure 5e). While the predicted preference varied across target contexts, the His6-tag with the highest predicted insertion rate was within the top two measured rates in each editing context (Supplementary Figure 6).

## DISCUSSION

We present the first comprehensive analysis of prime editing insertion efficiencies using 2,666 pegRNAs. We discovered that short sequences insert with predictable frequencies across cell lines, target sites, and prime editor systems based on their length, GC content and tendency to form secondary structure. Together, these features enable it to accurately predict variation in insertion rates, and to choose optimal reagents for writing short stretches of DNA into genomes.

We uncovered a complex relation between insertion sequence length and efficiency. Sequences between 15 and 21 nt generally inserted well, while longer sequences are incorporated less frequently, but still at moderate efficiencies even for sequences larger than 60 nt. This disparity is potentially due to steric issues for reverse transcription and base pairing with the unedited strand. The insertion efficiency of sequences shorter than 10 nt was variable, with high rates in MMR-deficient cell lines (HEK293T, HAP1 ΔMLH1) but not in MMR-proficient ones (HAP1). This is consistent with recent findings that MMR antagonizes prime editing^17^. Longer sequences are less efficiently recognized by MMR^30^ and therefore insert better than short ones in mismatch repair proficient cell lines.

We further discovered that stronger secondary structure of the pegRNA 3’-extension led to higher insertion efficiency. This effect was evident when comparing different inserts into the same target, but also explained variable rates when attempting to write the same sequence into different target sites. One potential explanation is that structured pegRNAs are more protected from digestion by 3’-exonucleases. Indeed, Nelson and Randolph *et. al*. recently demonstrated that incorporating structured motifs at the 3’-end of pegRNAs improved prime editing efficiency 3 to 4-fold by preventing degradation of the 3’-extension^18^. Alternatively, structured inserts could ease pairing of the edited strand with the non-edited strand due to being sterically smaller via folding onto themselves.

Our improved understanding of insertion efficiency using the prime editing system naturally leads to recommendations for experimental design. First, we suggest choosing sequences with high GC, and especially cytosine content that are prone to form secondary structures. Sequences whose lengths vary between 15 and 21 nt are well inserted using prime editing while sequences between 10 and 14 nt are not. Knocking out MLH1 will drastically improve insertion rates for sequences shorter than 10 nt. To boost the efficiency of short sequence insertions in mismatch repair proficient contexts, additional silent mutations should be inserted on the reverse transcriptase template or mismatch repair could be transiently inhibited (as implemented in PE4 or PE5 systems)^17^.

These factors can influence the choice of tag and codon. For example, the His-6 tag, especially if choosing the CAC codon, inserts almost 6 times as well as the next best tag in our library (Myc-tag). For the correction of pathogenic deletions, our model can help prioritize targets and pick high efficiency replacement sequences (for example through codon variation). Our libraries cover many commonly used small sequences and their respective insertion efficiencies (Supplementary Data 2). For predicting the insertion efficiency of novel sequences, we provide the MinsePIE algorithm as a command line script.

We studied four target sites in two cell lines across three prime editor systems and uncovered both universal and target site-dependent determinants of insertion efficiency. Models trained specifically on one target site still outperformed predictions on withheld data from the same target site when compared to predictions on other target sites. Screening a smaller focussed library across many target sites should help to more fully understand the interactions of target site and insertion sequences. Moreover, all pegRNAs had a constant primer binding site length of 13 nt and a reverse transcriptase template length (without insertion) of 34 nt. Given the model based on our sequence features (that implicitly also quantify these factors) generalised well to unseen pegRNAs, we expect the determinants we uncovered to also broadly hold for different lengths of primer binding site and reverse transcriptase template.

The prime editing field is moving rapidly^38^. Diverse applications are already emerging^39^ and some of the most exciting ones are specifically built around insertion of short sequences. Communicated, but as yet unpublished examples include insertion of recombinase sites using prime editing to enable directed insertion of large DNA cargo of up to 36 kb^1,2^, as well as clever utilization of short sequence insertion to generate a molecular recorder for sequential cellular events^40-42^. Better understanding of how cellular determinants^17^ and pegRNA features affect prime editing rates^14,18^ provides a foundation for these advances. Our work adds the important dimension of short sequences insertions, which hold the promise to both enable sophisticated genome engineering and to correct thousands of pathogenic mutations.

## Supporting information

Table S1 Primers

Table S2 pegRNAs

Table S3 Gene fragments

Data file 1 Insert sequence library

Data file 2 Insert sequence frequencies

Data file 3 Oligonucleotide pool

## Author contributions

Conceptualized and initiated the study: JK, LP. Performed experiments: EMP with help from: JK, FL. Analysed the data: JK, JW with help from: AP. Build the machine learning models: JW with help from: AP. Supervised the project: LP with help from: FL. Wrote the manuscript: JK, JW, LP with input from all authors.

## Competing interests

The authors do not declare any competing interests

## Acknowledgements

We thank Balca R. Mardin and Özdemirhan Serçin for discussions on the discrepancies in insert rates for short sequences between HEK293T and HAP1 cell lines, and for pointing us towards the mismatch repair pathway; We thank Thomas Vanderstichele, Matthew Coehlo, Erica Bello, Jacob Hepkema, Megan Gozzard, Luca Crepaldi and Tom Ellis for discussions and input on the manuscript. We thank Alexander Klenov for sharing the sequence of the MashUp Reverse Transcriptase derived from Feline Leukaemia Virus (FeLV). AP, EMP, JK, JW, and LP were supported by Wellcome (206194). LP was also supported by the Estonian Centre of Excellence in IT (EXCITE) (TK148).

## METHODS

### Mammalian cell culture

The human HEK293T cell line was purchased from AMS Biotechnology (EP-CL-0005). The HAP1 WT cell line was provided by Andrew Waters (Wellcome Sanger Institute) and the HAP1 ΔMLH1 cell line was purchased from Horizon discovery (HZGHC000343c022). HEK293T cells were cultured in DMEM (Invitrogen) and HAP1 cells in IMDM (Invitrogen), both supplemented with 10% FCS (Invitrogen), 2 mM glutamine (Invitrogen), 100 U/ml penicillin and 100 mg/ml streptomycin (Invitrogen) at 37 °C and 5% CO2.

### Primers

All primers used in this study are listed in Supplementary Table 1.

### Library design

The insert sequence library contained 2,666 sequences, made up from useful molecular biology sequences, the eukaryotic motif library (ELM) and sequences with strong secondary structure. The utility sequences were hand-picked for their usefulness in molecular biology. The ELM instances library with the corresponding fasta file of the genes was downloaded from elm.eu.org/instances.htmlãq=*^26,27^ on 2020/11/19 and filtered to only contain sequences from “homo sapiens” that are longer than 1 amino acid. The amino acid motifs were extracted from the fasta file based on the indicated start and end sites. Finally, the amino acid motifs were reverse translated into DNA sequence using the ‘reversetranslate’ R package (version 1.0.0) and using the most frequent codon from the homo sapiens codon table. For the secondary structure library, 100,000 random DNA sequences of 20 and 30 nt length were generated (RBioinf::randDNA function; version 1.48.0) and their secondary structure calculated (see insert sequence structure section). The sequences were distributed into 10 bins based on the strength of their secondary structure and 20 sequences were randomly picked from each structure bin to be included in the library. Finally, 30 random perfect 20 and 30 nt RNA hairpins were generated and amended to the secondary structure library. The combined library of insert sequences is included as Supplementary Data 1. The insert sequences were then flanked with primer binding sites, random nucleotide stuffer sequence for shorter inserts, BsmBI sites and target vector compatible overhangs, resulting in 11,166 sequences of 199 nt (Supplementary Data 3). The oligonucleotide library was ordered from Twist Biosciences.

### Plasmid cloning

*pCMV-PE2-P2A-PuroR* was generated by replacing eGFP from pCMV-PE2-P2A-GFP (Addgene 132776) with PuroR. Therefore, a gene fragment containing parts of the MMLV reverse transcriptase and the puromycin resistance gene was ordered from IDT (Supplementary Table 3). The gene fragment and pCMV-PE2-P2A-GFP were digested using AgeI, purified with the Monarch PCR & DNA Cleanup Kit (NEB) and ligated with T4 DNA ligase (NEB). 2 μl of the ligation product was transformed into bacteria using XL10-Gold Ultracompetent Cells (Agilent) according to the manufacturer’s protocol. Plasmid DNA was isolated using the Plasmid Plus Midi Kit (Qiagen).

*pCMV-PE-FeLV-P2A-EGFP* was generated by replacing the MMLV coding sequence between the XTEN linker and the 2A cleavage peptide with a synthesised gene fragment from IDT using Gibson Assembly that encodes an IDT human codon optimised version of the MashUp reverse transcriptase (pipettejockey.com) that is engineered from the Feline Leukaemia Virus (UniProt Q85521).

*pLentiGuide-Blast* was generated by replacing the puromycin resistance gene from Lenti_gRNA-Puro (Addgene 84752) with a blasticidin resistance gene. A gene fragment containing parts of the EF1a promoter and the blasticidin resistance gene was ordered from Twist Biosciences (Supplementary Table 3). The gene fragment and Lenti_gRNA-Puro were digested using FseI (NEB) and MluI-HF (NEB), purified with the Monarch PCR & DNA Cleanup Kit (NEB) and ligated with T4 DNA ligase (NEB). 2 μl of the ligation product was transformed into bacteria using XL10-Gold Ultracompetent Cells (Agilent) according to the manufacturer’s protocol. Plasmid DNA was isolated using the Qiagen Spin Miniprep Kit.

### Library cloning

First, a separate, site-specific backbone was cloned for each target site. A gene fragment was ordered containing the protospacer, guide RNA scaffold, parts of the reverse transcriptase template and primer binding site, a stuffer sequence flanked with BsmBI sites for insert library insertion and the T7 terminator motif (Supplementary table 3). 100 ng of the gene fragments were digested with BsaI-HFv2 (NEB) and purified with the Monarch PCR & DNA Cleanup Kit (NEB). The pLentiGuide-Blast plasmid was digested with BsmBI-V2 (NEB) at 55°C for 8h followed by 20 min heat inactivation at 80°C and gel purified using the QIAEX II Gel Extraction Kit (Qiagen). The gene fragments were ligated into the backbone using T4 DNA ligase (NEB) and transformed into XL10-Gold Ultracompetent bacteria (Agilent). The plasmids were purified with Qiagen Spin Miniprep Kit.

Second, pegRNA insert libraries were inserted into the site-specific backbones. The insert libraries were synthesized as 199 nt oligo pools (Twist BioSciences) and amplified using KAPA HiFi HotStart ReadyMix (Roche). Libraries for individual target sites were amplified with separate primers (Supplementary Table 1). The products were purified using the Monarch PCR & DNA Cleanup Kit, digested with BsmBI-v2 at 55°C for 4h and heat inactivated at 80°C for 20 min alongside 5 μg of site-specific plasmids. The digested oligos were purified using the Monarch PCR & DNA Cleanup Kit. The vectors were treated with quick CIP (NEB) for 15 minutes at 37°C and then purified using QIAquick PCR Purification Kit (Qiagen). Inserts were ligated into vectors using Golden Gate assembly. A 1:3 molar ratio of insert and vector were mixed with BsmBI-v2 and T4 DNA ligase and incubated in a thermocycler for 30 cycles, alternating between 5 minutes at 42°C and 5 min at 16°C and finishing with a heat inactivation step at 60°C for 5 min. The ligation products were purified with Monarch PCR & DNA Cleanup Kit and electroporated into MegaX DH10B T1R Electrocomp Cells (Thermo Fisher) according to manufacturer’s protocol. The bacteria were grown overnight in liquid culture and plasmid was extracted using the Plasmid Plus Midi Kit. The spacer sequences, primer binding sites, and reverse transcriptase templates (without insertions) are attached as Supplementary Table 2).

### Lentivirus production

Lentivirus was produced in HEK293FT cells that were transfected with Lipofectamine LTX (Invitrogen). 5.4 μg of a lentiviral vector, 5.4 μg of psPax2 (Addgene 12260), 1.2 μg of pMD2.G (Addgene 12259) were mixed in 3 ml Opti-MEM together with 12 μl PLUS reagent and incubated for 5 min at room temperature. 36 μl of the LTX reagent was added and the mix was incubated for another 30 min at room temperature. 3 ml of the transfection mix was then added to 80% confluent cells in 10 ml DMEM media in a 10-cm dish. After 48h the supernatant was collected and stored at 4°C. Fresh media was added to the cells and harvested 24h later. The two harvests were kept separate. For virus titration, Lenti-X GoStix Plus (Takara) was used following the manufacturer’s protocol.

### pegRNA insertion screens

*Infection with pegRNA library*. All cell lines were infected with the pegRNA library aiming at a multiplicity of infection (MOI) of 0.5 and a guide coverage of > 1000x. Each screen was performed in 3 biological replicates and independently infected. To achieve this, 6×10^6^ cells were plated in three wells of a six well plate and spin infected for 15-30 mins at 2000 rpm. Following infection, cells were resuspended and replated at 2×10^4^ cells/cm^2^. Cells were cultured for 7 days and selected for pegRNA integration with 10 µg/ml blasticidin. Cells were passaged at day 3 post-infection and a higher coverage than at the time of infection was maintained.

*Transfection with prime editors*. HEK293T cells were seeded at a concentration 6.9×10^4^ cells/cm^2^ to a 15-cm dish. The next day the media was replaced with fresh media and the cells were transfected using Lipofectamine LTX reagent. Next, 72 µg PE-Puro or PE-FeLV plasmid were mixed with 8 µg pCS2-GFP and 40 µl Lipofectamine P3000 (Invitrogen) in 3.2 ml Opti-Mem (Gibco). In another tube, 40 µl of Lipofectamine 3000 and 160 µl Lipofectamine LTX were mixed in 3.2 ml Opti-Mem. The solutions were mixed together, incubated for 30 minutes at room temperature and then added onto the cells. For PE3, an additional 6 µg of nicking guide RNA was added.

Hap1 and HAP1 ΔMLH1 cells were seeded at a concentration of 9.3×10^4^ cells/cm^2^ into two T75-flasks. The following day, the media was refreshed one hour before transfection. Cells were transfected using Xfect Transfection Reagent (Takara). Next, 72 µg PE-Puro plasmid was mixed with 8 µg pCS2-GFP and Xfect Reaction Buffer to a total volume of 750 µl. To the reaction, 48 µl of l Xfect Polymer was added and incubated at room temperature for 10 minutes. The mixture was then added onto the cells. Media was changed 4 h after transfection. One day after transfections, 2 µg/ml of puromycin was added to the cells to start selection. Cells were kept in selection for 3 days and harvested 5 days after transfection.

### DNA extraction and library preparation for next generation sequencing

Genomic DNA extraction and sequencing library preparation for screens were done as described in Allen et al., 2019^10^. Briefly, cell pellets were resuspended in TAIL BUFFER A (100 mM Tris-HCl, 5 mM EDTA, 200 mM NaCl) and then mixed with 1 volume of TAIL BUFFER B (100 mM Tris-HCl, 5 mM EDTA, 200 mM NaCl, 0.4% SDS) supplemented with freshly thawed Proteinase K (20 mg/ml final). The lysate was incubated overnight at 56°C. On the next day, RNase A was added to a final concentration of 10 µg/ml and incubated at 37°C for 30 min - 4 h. 1 volume of isopropanol was added and the DNA spooled on a sterile inoculation loop. The DNA was washed three times by dipping into consecutive 5 ml tubes containing 70% ethanol. The DNA was air dried for 5-10 mins and resuspended in TE buffer (pH 8.0).

For each screen, two independent amplicons were generated by PCR using Q5 Hot Start High-Fidelity 2X Master Mix (NEB). One amplicon for the targeted locus and one amplicon of the pegRNA locus (primers listed in Supplementary Table 1, staggered forward primers were used to ensure complexity for sequencing). To ensure coverage for each sample, 40 μg of gDNA was used as template and each PCR reaction was run in 50 μl aliquots containing no more than 5 μg DNA. The PCR reactions were column-purified using the QIAquick PCR Purification Kit (Qiagen). Sequencing adaptors and barcodes were added with a second round of PCR using the KAPA HiFi HotStart ReadyMix (Roche), primers P3 and P4 (Supplementary Table 1) and 1 ng of template DNA. Amplicons were purified with Agencourt AMPure XP beads in 0.7:1 ratio (beads to PCR reaction volume) and quantified with the Quant-iT™ High-Sensitivity dsDNA Assay Kit (Invitrogen). The amplicons were pooled together and sequenced on the Illumina HiSeq 2500 using HiSeq Rapid SBS Kit v2 (500 cycles, no phiX addition).

### Generating read count tables

Paired forward and reverse reads from illumina sequencing were merged using PEAR v0.9.11. Data for the same screen but from different sequencing lanes was concatenated. The resulting merged fastq files were processed using a custom R script (read_match_pegRNAs.R in Supplementary Information). First, DNA sequences were trimmed to contain the 10 nt up and downstream of the nick site (for target site amplicon) or to contain 15 nt up and downstream of the nick site (pegRNA amplicon). On average, 98% of reads were matched for the target site amplicon and 84% for the pegRNA amplicon. The trimmed sequences were then matched to each insert in the pegRNA library in the context of 10 nt target site (for target site amplicon) or in the context of 15 nt pegRNA plasmid (pegRNA amplicon), requiring 0 mismatches. Adding the context is to ensure that only insertions at the correct location are considered. On average 92% of reads were matched to the unedited locus or an insertion for both the target site amplicon and the pegRNA amplicon. The read count tables are attached as Supplementary Data 2.

### Combining replicates

We filtered out pegRNAs where any replicate had fewer than 10 reads in the pegRNA amplicon mapping to it. Across the screens, between 39 and 174 pegRNAs did not pass this minimum requirement, and were discarded from further analysis (1.5-6.5%, Supplementary Figure 1d). pegRNA abundance in the screens correlated with their abundance in the plasmid library (range of Pearson’s R across all samples 0.84 to 0.96, Supplementary Figure 1e). Insert counts were normalized to frequencies by dividing the reads for each insert by the number of reads in each screen. Insertion efficiencies were calculated for each replicate and screen by dividing the target insert frequency by the pegRNA insert frequency. Finally, insertion efficiencies were averaged across replicates. The script used to combine replicates is attached in the supplementary information as ‘combine_replicates.R’. Insertion efficiencies were normalized (z-score) between screens and replicates by subtracting the corresponding mean insertion efficiency from each individual insertion efficiency and dividing it by the standard deviation of the insertion efficiency.

### Mutation rates around the insertion site and indel detection

The fastq reads of the target sites were trimmed by matching a stretch of ten nucleotides 60 nt up and downstream of the nicking site (CLYBL: TAGGGCTGGA, CAGAGTTCCA; EMX1: GAGGACAAAG, ATGGGGAGGA; FANCF: GTCTCCAAGG, AGCACCTGGG; HEK3: CTTTTTTTCT, AGCTTTTCCT). Occurrence of library insertions was detected by pattern matching the trimmed reads for library sequences. Going from outside to inside (with the nicking site being between the two innermost nucleotides), the occurrence of the four nucleotides was counted at every position. There is a non-reference SNP (G>A) in HEK293T cells for 2 of 3 alleles at position +9. The RT template on the pegRNA corresponds to the sequence of the minor allele (A). For indel detection, the trimmed reads were filtered in a series of steps. First, sequences with insertions at the nick site that perfectly match a sequence in the insert libraries were removed (this also means that our method cannot detect single/double/triple nucleotide insertions at the nick site because our library contains all possible singlets/doublets/triplets). Second, sequences which contained ‘N’ were removed. Third, sequences with a perfectly preserved sequence around the cut site were removed. Fourth, sequences that are 120 nt long were removed (120 nt corresponds to the length of a sequence without indels). The remaining sequences were classified as indels. The scripts used to call mutation rates and indels are attached in the Supplementary Information as ‘find_SNVs.R’ and ‘find_indels.R’.

### Data analysis

Length residuals were calculated by dividing sequences into length bins and dividing the insertion rate by the median insertion rate across the length bin. The length bins consisted of sequences from 1-4, 5-9, 10-14, 30-39, 40-49, 50-59, and 60-69. The sequences with lengths above 30 nt were divided into length bins of 10 nt because there were fewer longer sequences in the library. Melting temperature for the insert sequence was calculated using SeqUtils.MeltingTemp.Tm_NN from biopython. The Vienna fold (VF) algorithm^31,32^ was used to calculate the tendency of insert sequences to form secondary structures. RNA fold (version 2.4.16) was run on the insert sequences alone or on the insert sequences in context of the 13 nt PBS and 34 nt RT template with the --noPS parameter.

### Modelling

Categorical features were one-hot encoded. Scikit-learn models were applied using default parameters, if not stated otherwise. Lasso regression was performed with alpha = 0.1; Ridge regression was performed with alpha of 0.03 and Stochastic Average Gradient descent; Random forest had a maximum depth of 4 and 100 estimators; Multilayer perceptron regressor with 100 hidden layers was trained with 500 maximum iterations at a constant learning rate of 0.001 and ‘lbfgs’ solver. Gradient boosted tree from XGBoost^36^ was trained with a minimum loss reduction of 0.1, 100 trees, a learning rate of 0.1, maximum depth of 5, no L1 regularization on weights, 0.375 L2 regularization on weights, and a 0.95 subsample ratio of columns when constructing each tree. For training, unique insert sequences were split randomly into training and test sequences at a ratio of 0.8. Measurements for different target sites and cell lines were assigned into training and test data based on the grouping of insert sequences. For 10-fold cross validation, insert sequences were split randomly for every fold validation. The model was trained and predictions were evaluated using Pearson’s R based on the correlation between test data and corresponding predictions. SHapley Additive exPlanations (SHAP) values for the model and feature importance for the prediction of specific outcomes were calculated using the SHAP TreeExplainer and explainerModel^37^.

### Statistics and reproducibility

The *n* numbers denoted in the figure legends refer to independent experiments that were separately infected with the pegRNA library. No statistical methods were used to predetermine sample size. The experiments were not randomized and the investigators were not blinded to allocation during experiments and outcome assessment.

### Data and material availability

Read count tables for all screens, mutation frequencies at each position, sequences with indels and scripts necessary to reproduce the analysis are attached in the Supplementary Information. The pCMV-PE2-P2A-PuroR and pLentiGuide_BlastR plasmids will be made available on AddGene. Scripts and models are made available on https://github.com/julianeweller/MinsePIE.

### Software

PEAR (0.9.11); Python (3.8.10); Python packages: Biopython (1.79), scikit-learn (0.24.2), scipy (1.5.3), shap (0.39.0), statannot (0.2.3), XGBoost (1.4.0); R (4.0.2); RNA fold (2.4.16); R packages: Broom (0.7.9), ggpointdensity (0.1.0), RBioinf (1.48.0), reversetranslate (1.0.0), ShortRead (1.46.0), spgs (1.0-3), Tidyverse (1.3.1), Viridis (0.6.1).

## Notes

https://github.com/julianeweller/MinsePIE

### Competing Interest Statement

The authors have declared no competing interest.

