## Supplementary information for "Predicting efficiency of writing short sequences into the genome using prime editing"

**Supplementary Figures:**

Supplementary Figure 1. Reproducibility of insertion efficiencies

Supplementary Figure 2. Prime insertion screen in MLH1 knockout HAP1 cells

Supplementary Figure 3. Nucleotide composition and secondary structure affect insertion efficiencies

Supplementary Figure 4. Additional classes of sequences that affect insertion rate

Supplementary Figure 5. Marginal effects of length, GC-content and structure on insertion rate is consistent between PE-FeLV, PE3 and PE2

Supplementary Figure 6. Evaluating the prediction accuracy of MinsePie

**Supplementary Tables:**

Supplementary Table 1. Primers used in this study

Supplementary Table 2. pegRNAs used in this study

Supplementary Table 3. Gene fragments

**Supplementary Data Files:**

Supplementary Data 1. Insert sequence library

Supplementary Data 2. Insert sequence frequencies (read count table)

Supplementary Data 3. Library oligopool as ordered from Twist Biosciences

**Supplementary Code:**

Scripts used to analyze screen data: https://github.com/julianeweller/MinsePIE

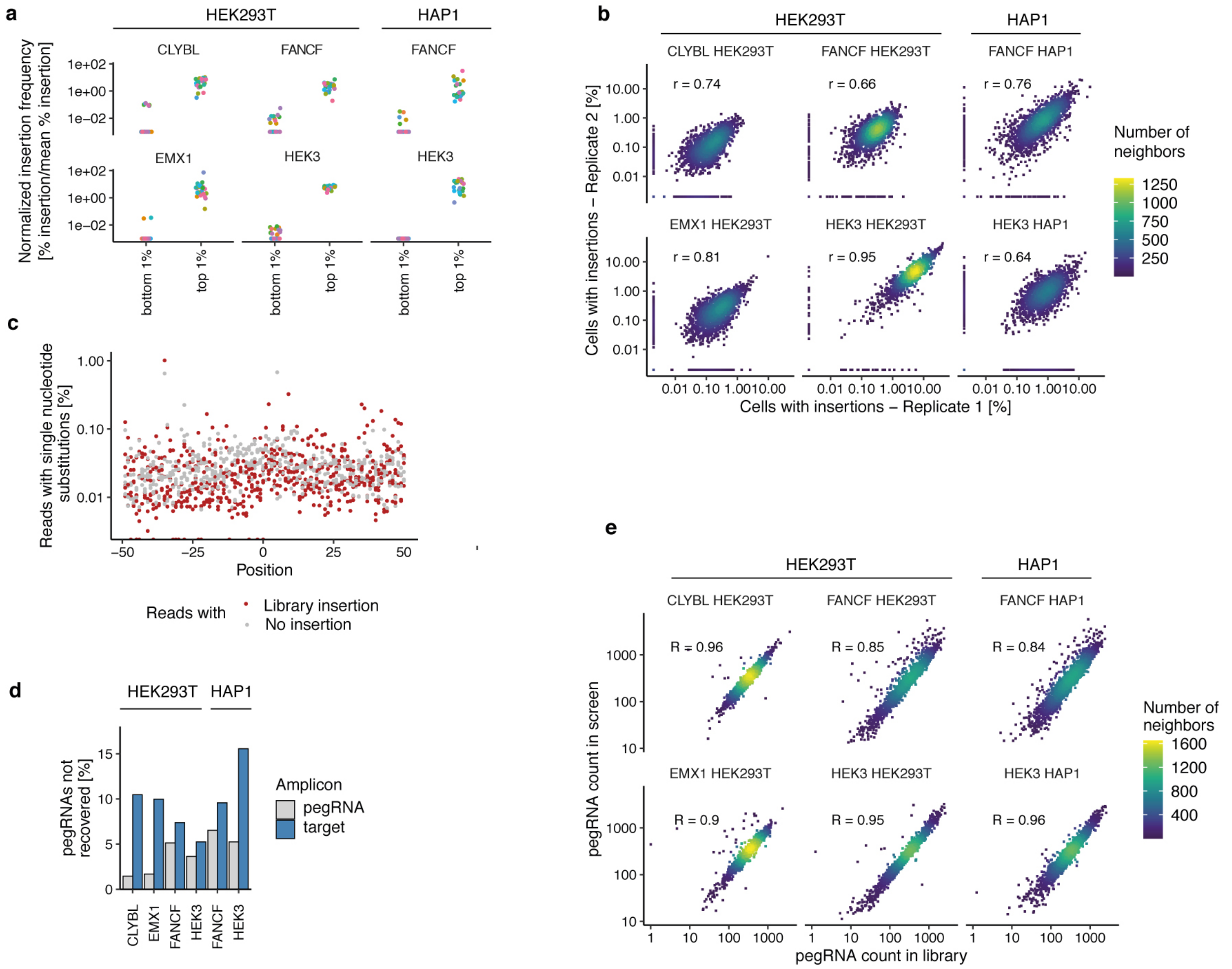


**Supplementary Figure 1. Reproducibility of insertion efficiencies. a.** Insertion efficiency of pegRNAs varies widely. Screen normalized insertion efficiency (y-axis) for the top 1% of pegRNAs with highest insertion rates across all screens and the bottom 1% of pegRNAs (x-axis). Markers are individual pegRNA colored by their identity. Data represent the average of *n*=3 biological replicates **b.** Insertion rates are highly reproducible. Percent insertion in replicate 1 (x-axis) compared to percent insertion in replicate 2 for insert sequences (markers) in different cell lines and target sites (panels). **c.** Single nucleotide mutations are rare. Average percentage of reads with a non-reference sequence nucleotide (y-axis) at positions relative to the nicking site (x-axis) for reads with no insertion (grey) and with an inserted sequence (red). *n*=3 biological replicates. **d.** Percentage of non-detected insert sequences (y-axis) for different screens (x-axis) stratified by amplicons (blue: target site amplicon; grey: pegRNA amplicon). **e.** pegRNA abundance in screen is highly correlated with pegRNA abundance in the plasmid library. pegRNA count in the plasmid library (x-axis) compared to pegRNA count in different screens (panels) for insert sequences (markers). Data represent the average of *n*=2 sequencing replicates (library) and *n*=3 biological replicates (screens).

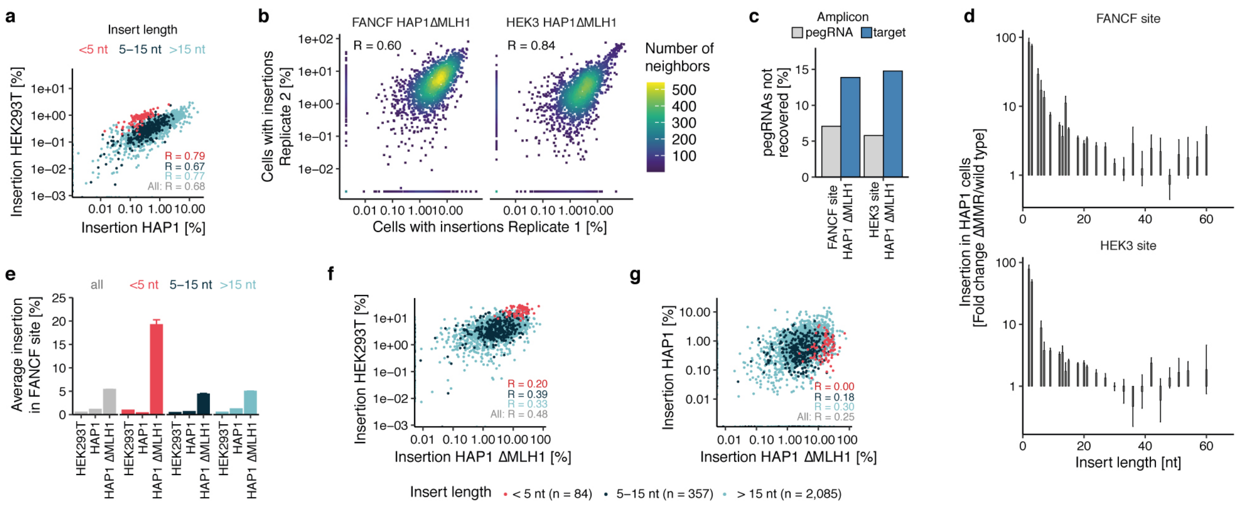


**Supplementary Figure 2. Prime insertion screen in MLH1 knockout HAP1 cells. a.** HAP1 cells insert short sequences at lower rates than HEK293T. Insertion rate in HEK293T cells (y-axis) compared to rate in HAP1 cells (x-axis) at the FANCF target of individual sequences (markers). Red: short sequences (up to 4nt); blue: medium sequences (5-15nt); teal: longer sequences (>15nt). Label: Pearson's R between rates. The data are an average from *n*=3 biological replicates. **b.** Replicate correlation for HEK3 and FANCF target sites in the HAP1 ΔMLH1 cell line. Percent insertion in replicate 1 (x-axis) compared to percent insertion in replicate 2 for insert sequences (markers) for FANCF target site (left panel) and HEK3 target site (right panel). **c.** Percentage of non-detected insert sequences (y-axis) for the FANCF and HEK3 sites (x-axis) stratified by amplicons (blue: target site amplicon; grey: pegRNA amplicon). **d.** MLH1 knockout increases insertion rates disproportionally for short sequences. Fold changes in insertion rates between HAP1 ΔMLH1 and HAP1 wild type cells (y-axis) for different insert sizes (x-axis) and target loci (panels). Bars: mean and s.e.m. Only insert sizes with more than 5 sequences are shown. **e.** Average insertion rates at the FANCF locus (y-axis) across different cellular contexts (x-axis) for different length bins (colors). Error bars: standard error of the mean. Colors as in (a). **f-g).** As (a), but comparing insertion rates in HAP1 ΔMLH1 cells with HEK293T cells (f) and HAP1 wild type cells (g).

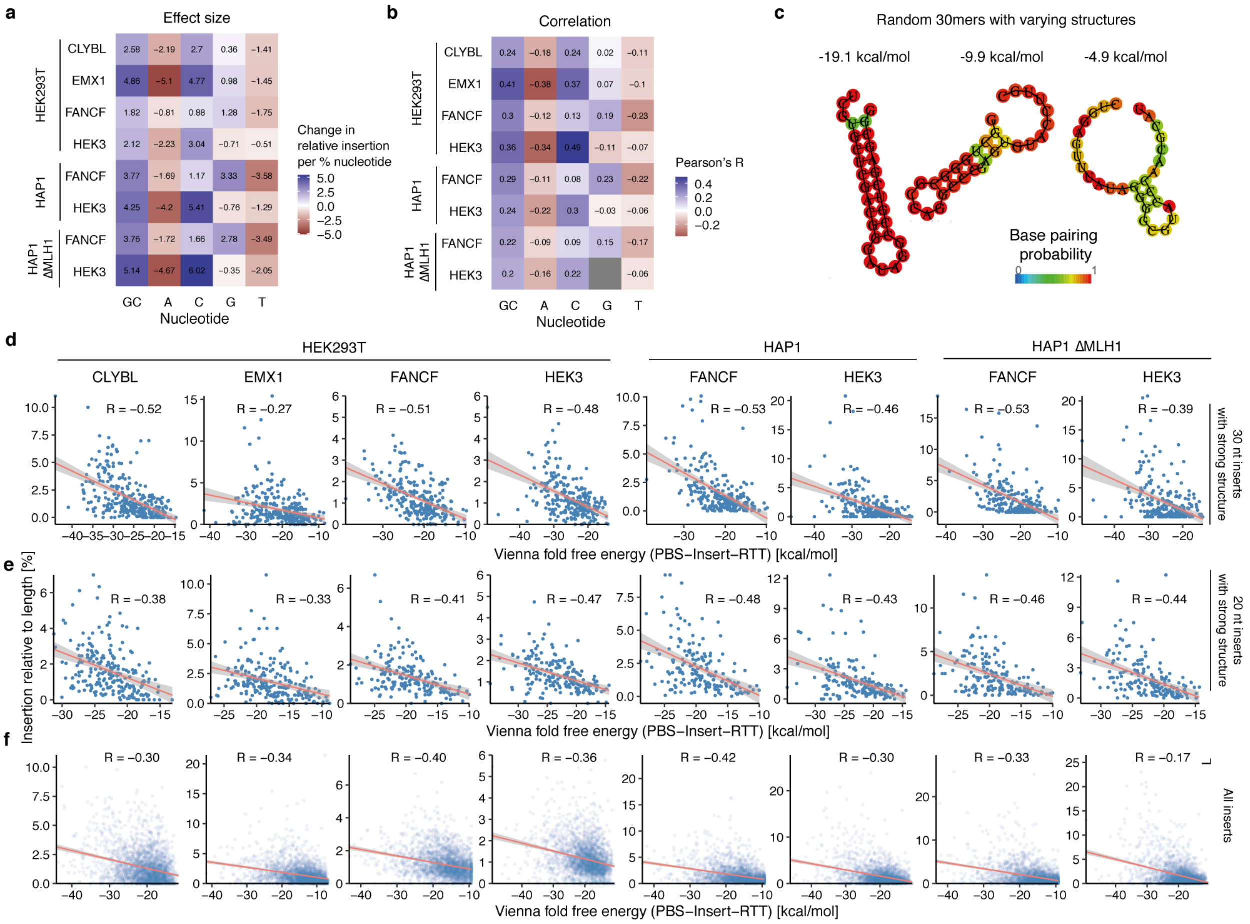


**Supplementary Figure 3. Nucleotide composition and secondary structure affect insertion efficiencies. a.** Percent additional insertion relative to the average length-normalized insertion rate (color) per extra percent of nucleotide in the insert sequence (x-axis) across different cell lines and targets (y-axis). Data represent the average of *n*=3 biological replicates. **b.** As (a) but instead displaying Pearson's R. **c.** Examples of predicted secondary structures and their corresponding free energies for three 30 nt sequences with differing structure strengths. Colors: base pairing probability. **d.** Length-normalized insertion rate (y-axis) for 30 nt insertions (markers) at different target sites and cell lines (panels) with calculated Gibbs free energy (ΔG) from ViennaFold (x-axis). Red line: linear regression fit; shaded area: 95% posterior confidence interval of the fit. Data represent the average of *n*=3 biological replicates. **e-f.** As in (d) but showing 20 nt insertions (e) or across inserts of all lengths (f).

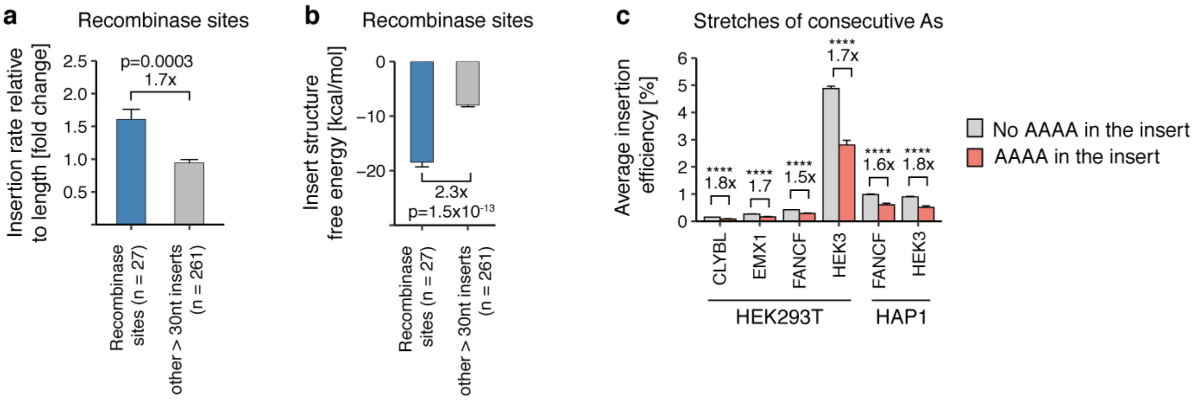


**Supplementary Figure 4. Additional classes of sequences that affect insertion rate. a.** Recombinase sites are efficiently inserted relative to their size. Average insertion rate relative to length bin (y-axis) for recombinase sites or other insert sequences larger than 30 nt (x-axis). Bars: Average and s.e.m. Comparison: ratio of blue to grey bar height. p-value: two-sided Student's t-test. *n*=3 biological replicates. **b.** As (a), but for the secondary structure of the inserts. **c.** Impact of four consecutive adenines. Average insertion rate (y-axis) for different screens (x-axis) stratified by presence of four consecutive adenines (grey: absent; red: present). Comparison: ratio of grey to red bar height. p-value: two-sided Student's t-test (all $<1x10^{-8}$). *n*=3 biological replicates.

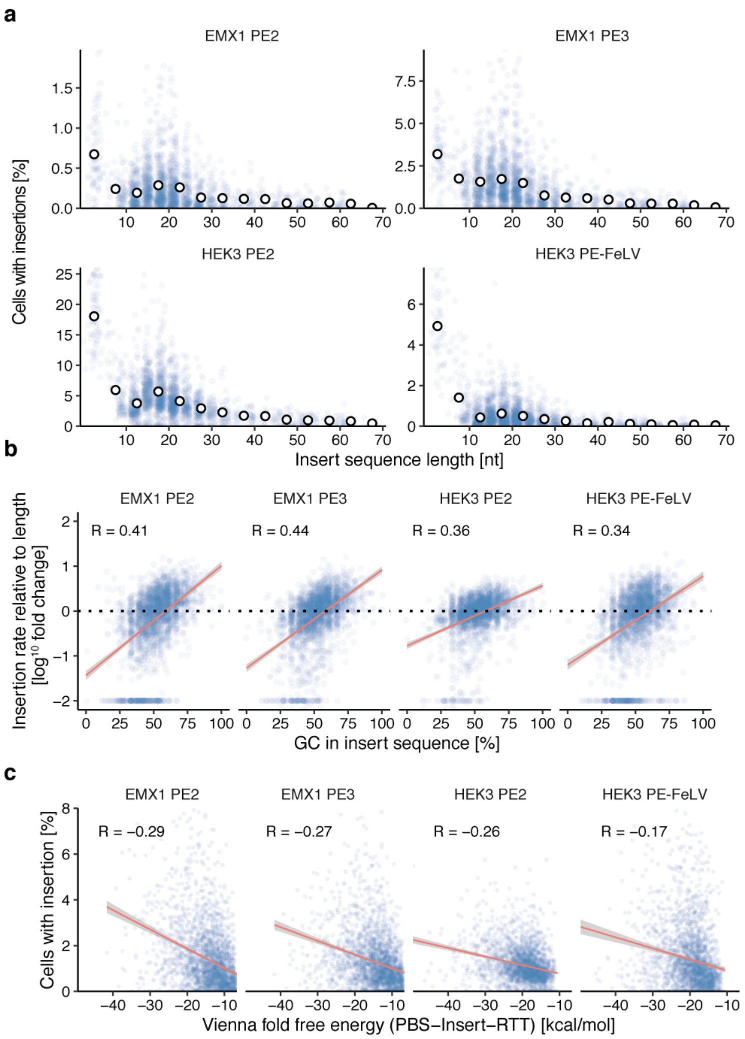


**Supplementary Figure 5. Marginal effects of length, GC-content and structure on insertion efficiency are consistent between PE-FeLV, PE3 and PE2. a.** Percent HEK293T cells with insertion (y-axis) for different insert sizes (x-axis) of individual sequences (blue markers) and their averages in 5nt length bins (white markers) at the EMX1 and HEK3 site using different prime editing reagents (panels). Data represent the average of *n*=3 biological replicates. **b.** GC content in the insert sequence increases insertion efficiency. log10-scale percent HEK293T cells with insertion relative to length bin average (y-axis) for inserts with different GC content (x-axis) of individual sequences (markers) at the EMX1 and HEK3 site using different prime editing reagents (panels). Red line: linear regression fit; shaded area: 95% posterior confidence interval of the fit. Data represent the average of *n*=3 biological replicates. **c.** Strength of insert sequence secondary structure is positively correlated with insertion rate. Percent of HEK293T cells (y-axis) with insertions (markers) at the EMX1 and HEK3 site using different prime editing reagents (panels) with calculated Gibbs free energy (ΔG) from ViennaFold (x-axis). Red line: linear regression fit; shaded area: 95% posterior confidence interval of the fit. Data represent the average of *n*=3 biological replicates.

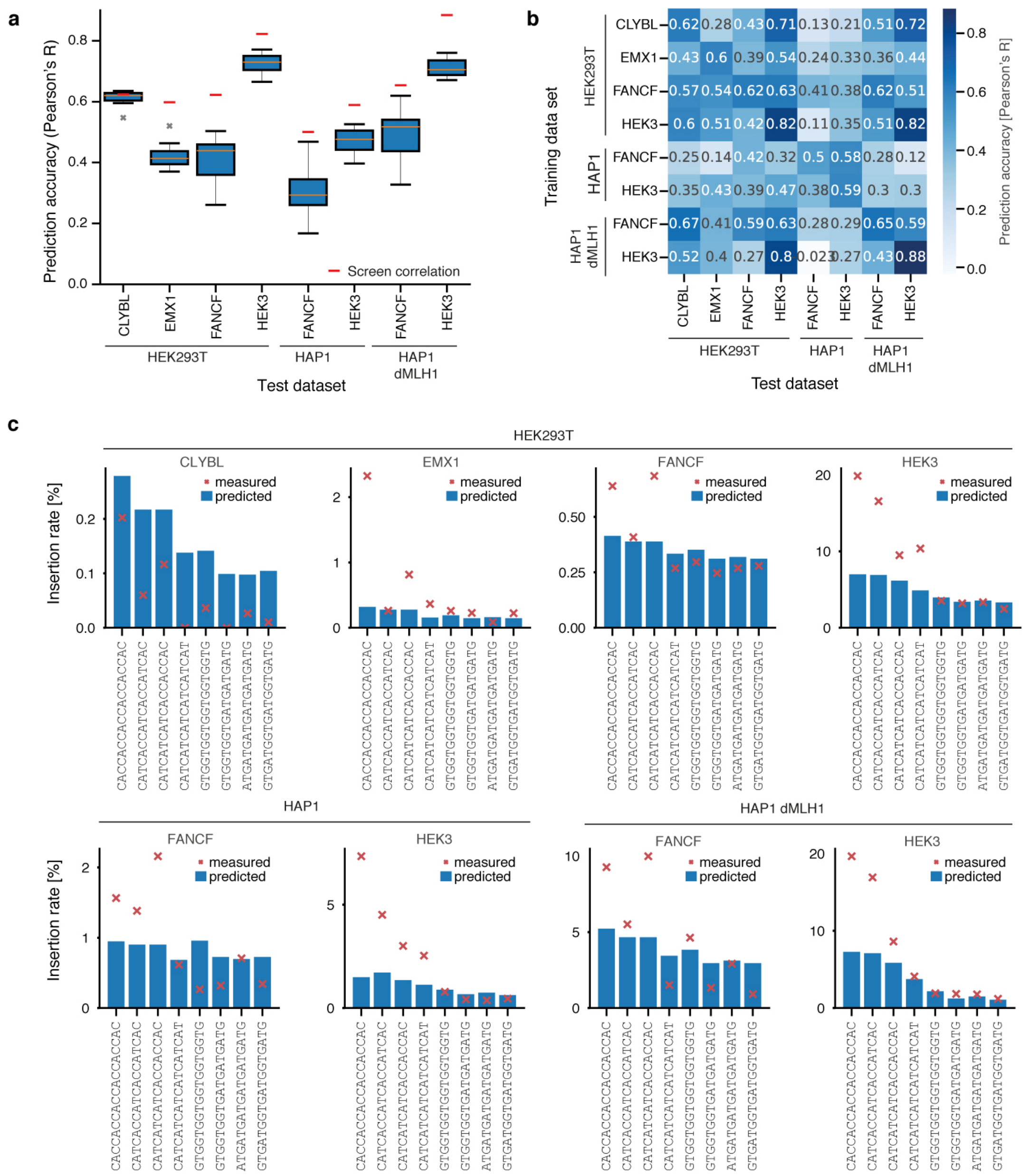


**Supplementary Figure 6. Evaluating the prediction accuracy of MinsePie. a.** Performance of XGBoost model on unseen target site in different cell lines. Pearson's R between predicted and measured held-out insertion sequences (y-axis) for ten cross-validation folds across a range of test datasets. For training, only datasets from other target sites were included. Box: median and quartiles; whiskers: range, x: outliers. **b.** Pairwise Pearson's R between predicted and measured held-out insertion sequences of MinsePIE trained on a single editing context (y-axis) and tested on another context (x-axis). **c.** Comparison of predicted (blue bars) and measured (red cross) insertion rates for His6-tags across target sites and cell lines.
